# No Strings Attached: Predicting Tricuspid Valve Deformation Without In Vivo Chordal Geometry

**DOI:** 10.64898/2026.09.25.754571

**Authors:** Mrudang Mathur, Collin E. Haese, Vijay K. Dubey, Sumedh Seetharam, William D. Meador, Tomasz Jazwiec, Natalie T. Simonian, Michael S. Sacks, Marcin Malinowski, Jan Fuhg, Issam Moussa, Joshua A. Cohen, Matthew R. Summers, Tomasz A. Timek, William Hiesinger, Manuel K. Rausch

**Affiliations:** Walker Department of Mechanical Engineering, The University of Texas at Austin, Austin, 78712, Texas, U.S.; Department of Cardiothoracic Surgery, Stanford University, Stanford, 94305, California, U.S.; Department of Biomedical Engineering, The University of Texas at Austin, Austin, 78712, Texas, U.S.; Department of Cardiac, Vascular and Endovascular Surgery and Transplantology, Silesian Centre for Heart Disease, Medical University of Silesia in Katowice, Zabrze, Poland; The Oden Institute of Computational Science and Engineering, The University of Texas at Austin, Austin, 78712, Texas, U.S.; Department of Cardiac Surgery, Wojewódzki Szpital im. Sw. Ojca Pio, Przemyśl, Poland; Department of Aerospace Engineering & Engineering Mechanics, The University of Texas at Austin, Austin, 78712, Texas, U.S.; Carle Illinois College of Medicine, University of Illinois at Urbana-Champaign, Urbana, 61801, Illinois, U.S.; Sentara Heart Valve Center, Sentara Heart Hospital, Norfolk, Virginia, U.S; Division of Cardiothoracic Surgery, Corewell Health West, Grand Rapids, 49503, Michigan, U.S.; College of Human Medicine, Michigan State University, Grand Rapids, 49503, Michigan, U.S.

**Keywords:** hyperelastic warping, image registration, ultrasound, transcatheter, repair

## Abstract

Predictive biomechanical models of the tricuspid valve require accurate representation of the chordae tendineae, yet subject-specific chordal geometry is difficult to reconstruct from non-invasive imaging. Here, we adopt a framework for generating functionally equivalent synthetic chordae without prior knowledge of in vivo chordal attachments. To this end, we first adapt an anatomy-informed hyperelastic shape-matching method that establishes correspondence between end-diastolic and end-systolic leaflet configurations using chordal-mimicking forces, a rigid contact template, and a Gaussian-smoothed locally corrective pressure field. We then generate synthetic chordal insertion sites using zone-based rejection sampling and calibrate the unloaded length of each chord by combining reaction forces with the chordal stress–stretch relationship. The framework was evaluated using Texas TriValve 1.1, a high-fidelity finite element model of a human tricuspid valve validated against beating-heart echocardiography. We found that shape matching reproduced the target end-systolic geometry with a mean inter-surface distance of **0.29 *±* 0.35** mm. Moreover, synthetic chordal insertions faithfully reproduce end-systolic leaflet deformations. We subsequently examined synthetic chordal configurations containing 202, 225, and 450 insertions, informed by measurements from eight explanted human tricuspid valves. Here, increasing insertion number reduced mean inter-surface distance from **0.63 *±* 0.52** mm to **0.49 *±* 0.44** mm and reduced contact area errors from 3.52% to 0.59%. Across all configurations, mean maximum principal stretch errors in leaflet belly regions remained below 2.4%, while areal strain errors ranged from 0.34% to 10.01%. These results demonstrate that anatomically informed shape matching coupled with stress-based chordal calibration can reproduce tricuspid valve closure without explicit subject-specific chordal geometry. This framework provides a foundation for generating synthetic subvalvular anatomy for future imaging-derived, predictive tricuspid valve models.

## 1 Introduction

Tricuspid valve regurgitation has been declared a “public health crisis” due to its historic undertreatment[1]. While recent clinical guidelines advocate for more aggressive and proactive repairs[2], the efficacy of surgical and interventional therapies is still lacking[3, 4]. This is partly due to an incomplete consideration of valvular mechanics in the healthy, diseased, and repaired tricuspid valve at the patient level. Current repair guidelines and expert testimony recommend a standardized repair approach that is largely agnostic to valvular morphology and mechanics[5, 6]. However, we have previously demonstrated in sheep that surgical repairs alter the mechanical state of the tricuspid valve[7, 8]. Furthermore, we have computationally demonstrated that valvular mechanics are sensitive to surgical annuloplasty parameters and warrant optimization at the patient level[9].

High-fidelity computer models, created from patient imaging, are promising tools for optimizing valve repairs[10, 11]. Unfortunately, the widespread application of such models in planning tricuspid valve repair remains limited. This may be attributed to two critical challenges linked to building imaging-based models of the tricuspid valve. Both challenges are associated with echocardiography, the current clinical standard of care used assess valvular function[12]. First, it is difficult to consistently image the thin tricuspid leaflets over the cardiac cycle[13]. As a result, segmented valve geometries at end-diastole and end-systole may not share a common material point correspondence, thus exacerbating challenges in model validation. Second, it is particularly challenging to reconstruct the complete subvalvular apparatus, consisting of tricuspid chordae tendineae and papillary muscles, based on echocardiography alone[14]. Chordae tendineae tether the tricuspid leaflets while preventing prolapse and are, thus, critical to its functioning[15].

To address the first challenge, strategies such as Large Deformation Diffeomorphic Metric Mapping (LDDM) have long been used to build material point correspondence for mitral and aortic heart valves. However, they rely on certain foundational assumptions. For example, they assume that valve leaflets are visible throughout the cardiac cycle and are, therefore, precluded from topological changes between the end-diastolic and end-systolic segmented shapes[16]. These assumptions may prove to be brittle in applications for the tricuspid valve, where the valve’s orientation and location, limited acoustic windows, and thinner leaflets frequently lead to signal dropout[17]. As an alternative strategy, hyperelastic warping or “shape matching” may also be considered[18]. Here, a deformable registration problem is regularized by the hyperelastic energy functional of a finite element problem to ensure consistency with large-deformation kinematics. This technique was introduced by Rabbitt et al.[19] to model the left ventricle and has since been adapted to model heart valves. In particular, Rego et al. first detailed these methods for in vivo, transesophageal echo (TEE) images of ovine mitral valves[20] and human aortic valves[21]. Unlike LDDM-based methods, hyperelastic shape matching is robust to incomplete or “clipped” segmentations, as demonstrated by Simonian et al. for human mitral valves imaged with TEE[22]. To address the second challenge for the mitral valve, Khalighi and colleagues used a topology optimization framework to convert complex chordal insertions into simplified, branchless structures for ovine valves imaged with computed tomography in an in vitro setting[23]. Notably, they demonstrated that the branchless insertions faithfully reproduce valvular deformations when compared to their in vivo counterparts and are thus “functionally equivalent”. Despite these advances in mitral valve modeling, similar progress is lacking for the tricuspid valve.

In this study, our overall objective is to overcome these challenges and predict tricuspid valve deformation without any a priori knowledge of in vivo chordal geometry. To this end, we first integrate a novel chordal force substitution algorithm with established hyperelastic warping methods to build a material correspondence between the end-diastolic and end-systolic states of the valve. We then use a stress-based calibration strategy to build synthetic chordae tendineae. Finally, we demonstrate these methods for Texas TriValve 1.1, a reverse-engineered, openly available model of a human tricuspid valve.

## 2 Methods

### High-fidelity Finite Element Model

In this study, we use a subject-specific, high-fidelity finite element model of the human tricuspid valve built using our Texas TriValve (reverse) engineering pipeline (see Figure 1). While the details of this pipeline are available in our previous publication [24], we provide a brief description here for completeness. The model was developed using a healthy human donor heart that was rejected for transplantation and placed in an organ preservation system (OCS, Transmedics, Andover, MA). After reperfusing the heart with the donor’s blood and restoring it to sinus rhythm, we recorded right-heart dynamics using sonomicrometry crystals (Sonometrics Inc, London, ON, Canada), transvalvular pressure using pressure transducers (PA4.5-X6; Konigsberg Instruments Inc., Pasadena, CA), and leaflet dynamics using echocardiography (Vivid S6, GE Healthcare, Chicago, IL). Next, we excised the valve complex for further characterization. We used photogrammetric digitization and a thickness gauge to assess leaflet geometry. To quantify the constitutive responses of the leaflets and chordae tendineae, we performed biaxial and uniaxial tensile tests, respectively. For constructing the in silico model, we began by non-rigidly transforming the digitized leaflets onto the annulus to attain a stress-free state. Subsequently, the sub-valvular apparatus was added based on ventricular images and existing literature. The leaflets were discretized using reduced-integration quadrilateral shell elements (S4R), while chordae tendineae were digitized as truss elements (T3D2). We assigned a uniform thickness to the shell elements based on in vitro characterization of valve leaflets. Moreover, we assigned leaflet-specific cross-sectional areas to the truss elements also based on in vitro measurements. After preparing the geometry and discretization, we applied Dirichlet boundary conditions informed by the sonomicrometry data from the same beating heart and imposed a pressure-gradient Neumann boundary condition on the valve leaflets. Finally, we simulated leaflet coaptation in Abaqus/Explicit Abaqus (v20.1, Dassault Syst‘emes, Providence, RI), completing the modeling workflow. Please note that model constitutive and simulation parameters are detailed in the openly available Abaqus input files, see **Data Availability** below.

**Fig 1.**
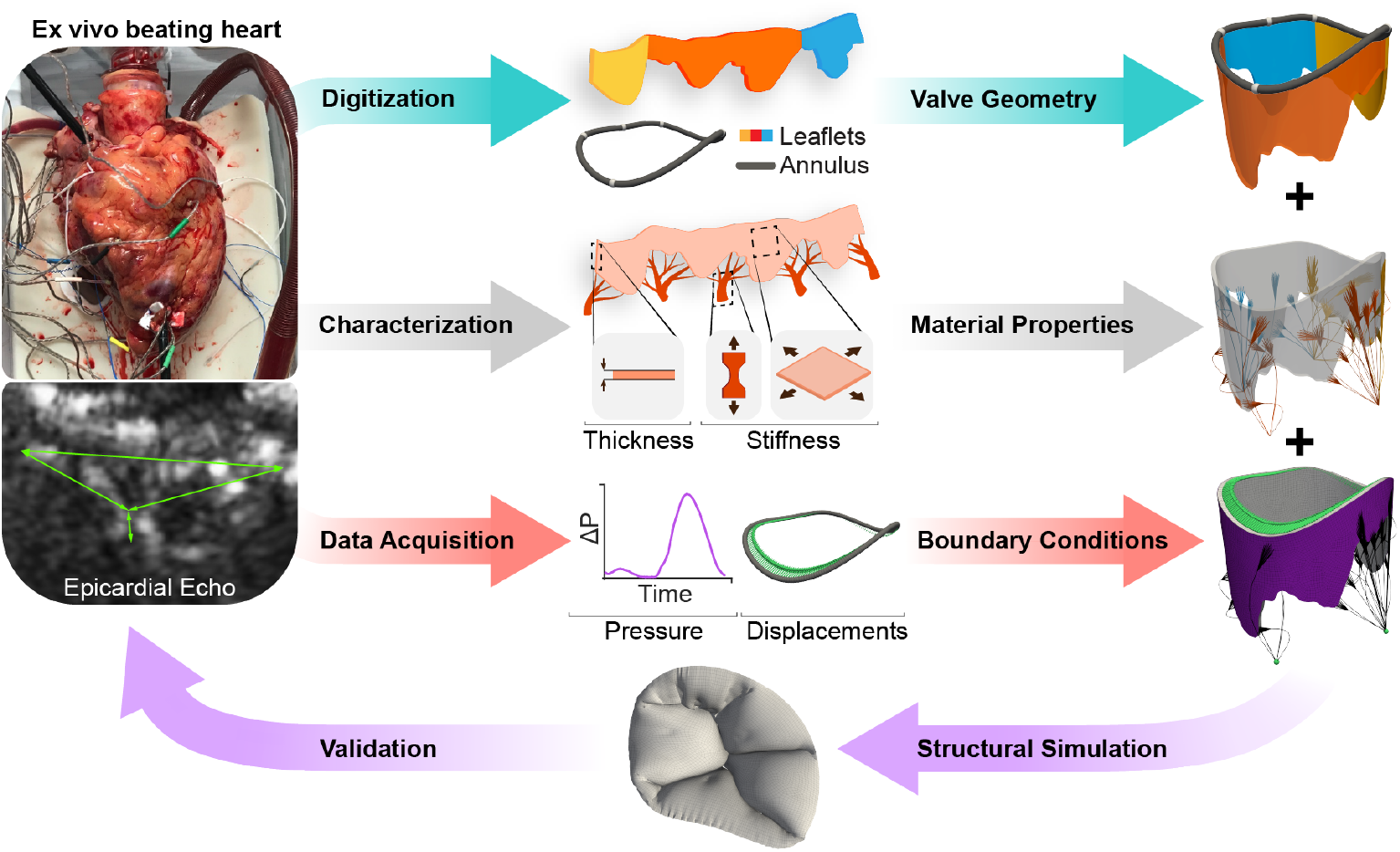
Texas TriValve modeling pipeline. We reverse-engineer a subject-specific model of the human tricuspid valve from an ex vivo, beating heart. Reproduced with permission from Mathur et al.[24]

### Hyperelastic Shape Matching

Image segmentations of the end-diastolic and end-systolic configurations of the tricuspid valve may not share the same triangulation and, thus, do not possess a natural correspondence between material points. To build such a correspondence, we rely on hyperelastic shape matching[25]. Like Rego and colleagues[20], we use a hyperelastic shape matching procedure determined by chordal mimicking forces and a spatially varying pressure field. We also introduce a rigid template to ensure correct inter-leaflet contact. The construction, calibration, and staged application of both are detailed in the following sections.

### Chordal Mimicking Forces

Chordae tendineae tether tricuspid leaflets to the ventricular myocardium and prevent their prolapse into the right atrium during systole. To that end, their apical or “downward”, action is well documented and understood. However, we posit that the chordae tendineae also tether the leaflets in the circumferential direction. Specifically, we have previously observed that tricuspid chordae tendineae insert into the leaflets at an angle from the radial direction[26]. To capture the overall effect of the tricuspid valve’s chordae tendineae we created two classes of chordal mimicking forces: downward forces and lateral forces. Both forces are applied to the free edge of the valve leaflets as demonstrated previously[20]. Simulation outcomes in the absence of lateral forces are provided in the **Supplementary Materials**.

For the downward forces, we generate unit vectors in the apical direction at each node along the free edge, see **Fig. 2A&B**. To assign the lateral forces, we first create a 2D projection of the leaflet surfaces by unwrapping the valve along the annulus; see **Fig. 2C**. On this projection, we split each leaflet cusp into two distinct sides — the left and right edges. The split locations are determined by a change in the sign of the tangent to the free edge curve on the 2D projection. Please note that we only split edges for cusps that exceed 3 mm in width to account for the inherent uncertainty in 3D transesophageal echocardiography measurements of atrioventricular valve leaflets[27]. We then designate each edge to repel a specific commissure, see **Fig. 2D**. To that end, the left-sided edges of the posterior leaflet repel the antero-posterior (AP) commissure while the right-sided edges repel the postero-septal (PS) commissure. Similarly, the left-sided edges of the anterior leaflet repel the antero-septal (AS) commissure while the right-sided edges repel the antero-posterior (PS) commissure. Finally, the left-sided edges of the septal leaflet repel the postero-septal (PS) commissure while the right-sided edges repel the antero-septal (AS) commissure. On every free-edge node, we then assign repulsive forces via.

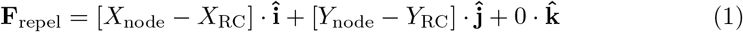

where {*X/Y*}_node_ represent the x-y Cartesian coordinates of each free-edge node in the reference configuration and {*X/Y*)_RC_ represent the x-y cartesian coordinates of the corresponding repelled commissure in the reference configuration. Finally, we normalize the repulsive force vectors before calibrating their magnitudes, see **Fig. 2E**.

**Fig 2.**
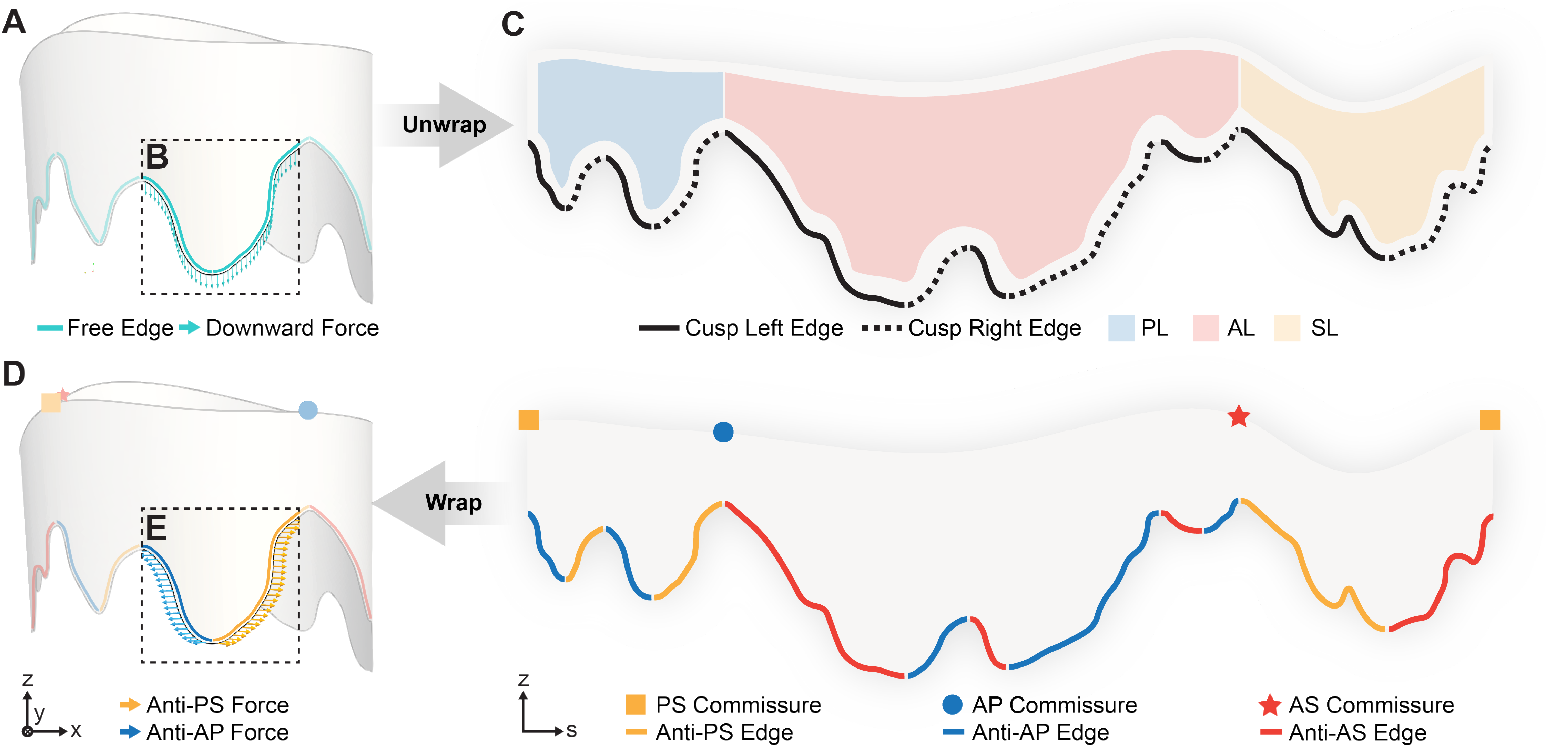
Generating Downward and Lateral Force Vectors. (A) We identify the free-edge of the valve and assign a (B, inset) downward force at each node. (C) We then “unwrap” the tricuspid leaflets along the circumferential direction of the valve annulus to create a 2D projection. On the 2D projection, we isolate the left and right edges of each major cusp. (D) Next, we mark each edge as anti-PS, anti-AP, or anti-AS to move away from one of two commissures located on either end of each leaflet. We then “wrap” the leaflet surface along the circumferential direction of the valve annulus to recreate the 3D valve. (E, inset) Finally, we generate lateral forces. AL: Anterior Leaflet, PL: Posterior Leaflet, SL: Septal Leaflet, AP: Antero-posterior, PS: Postero-septal, AS: Antero-septal

### Force Calibration and Rigid Inflation

Having generated both downward and lateral force vectors, our next objective is to calibrate their magnitudes. Following prior work on the mitral valve[20], we distribute a net force of specified magnitude, referred to as CMF_down_, uniformly across all leaflet free-edge nodes in the downward direction. In contrast, the lateral force vectors require careful treatment to ensure accurate leaflet coaptation. Unlike the mitral valve, where the anterior and posterior leaflets must contact each other, the multi-cusped nature of the tricuspid valve may lead to incorrect leaflet contact pairings. Thus, we must use a structured biasing strategy to control the motion of each cusp in the lateral direction, detailed in **Algorithm 1**. As a result, we reduce this bias calibration step to a simpler problem of tuning bias parameters {*α*_1_, *α*_2_, …, *α*_*k*_}, where *k* is the number of cusps within a given valve. We then distribute a net force of specified magnitude, referred to as CMF_lateral_, equally to each leaflet cusp in the lateral direction. Within each cusp, this force is weighted to one side of the leaflet and applied unequally to the free-edge nodes on each side. This results in a non-zero force vector in the lateral direction on each cusp, which “pulls” each leaflet cusp laterally.

Next, we created a rigid template of our target shape – i.e., the valve’s end-systolic configuration – and simulated leaflet coaptation in Abaqus/Explicit, see **Fig. 3A**. In Stage I, we applied a generic pressure gradient of 22 mmHg on the ventricular surface of the leaflets, applied both downward and lateral forces, and enabled contact between the rigid template and our valve model. We calibrated the magnitude of the downward and lateral total forces (CMF_down_ and CMF_lateral_, respectively) by minimizing the following error:

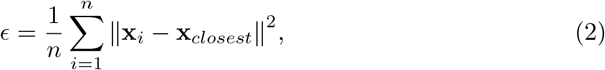

where *n* is the number of nodes, **x**_i_ is the nodal position of the predicted shape, and **x**_closest_ is the position of the closest point on the target shape. Further details on the optimization procedure are provided in the **Supplementary Materials**.

**Fig 3.**
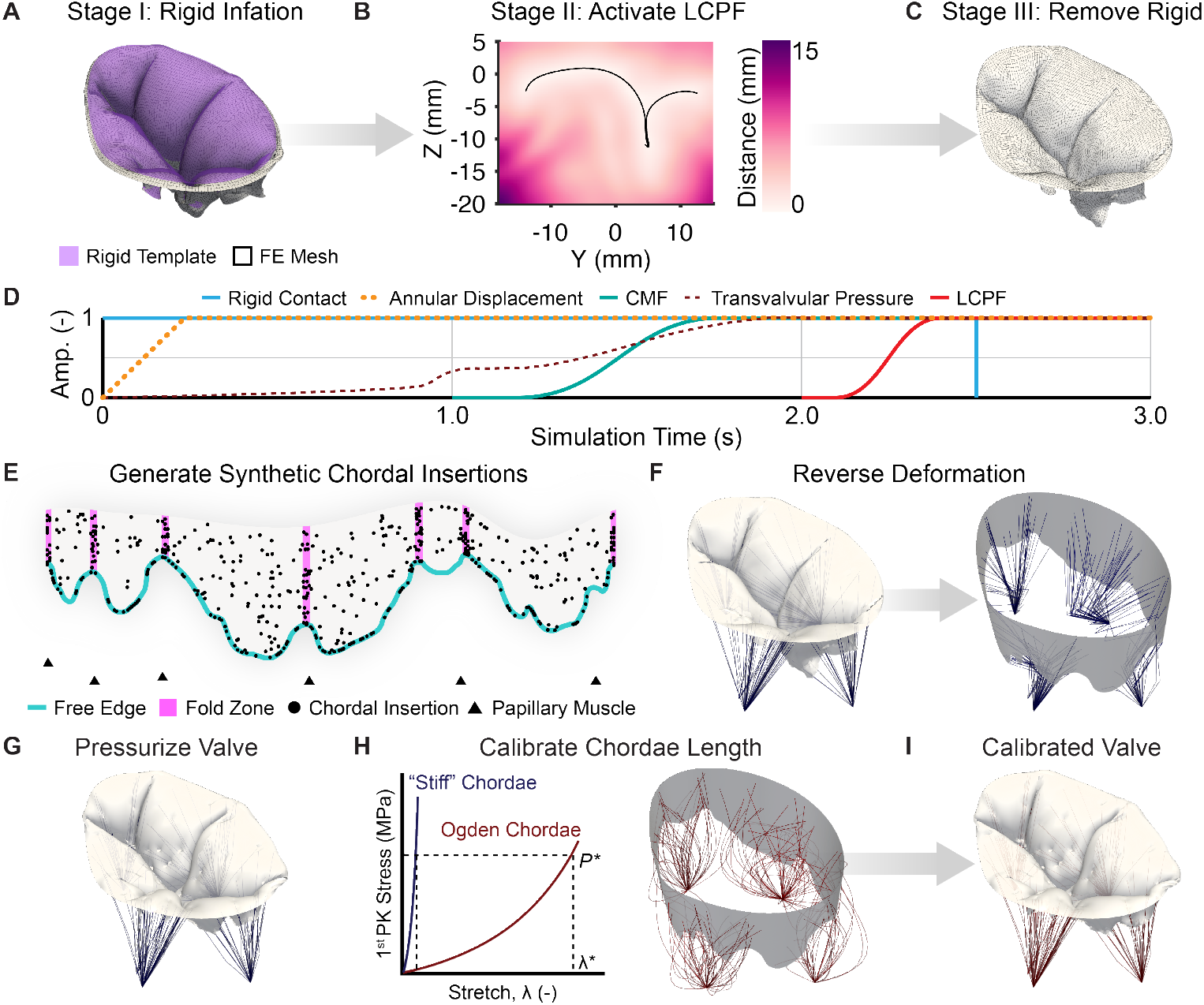
Shape matching and synthetic chordae generation pipeline. We perform hyperelastic shape matching in three stages: (A) Stage I – inflation against a rigid template with chordae mimicking forces (CMF), (B) Stage II – the addition of a locally corrective pressure field (LCPF), and (C) Stage III – removal of the rigid template and equilibration of forces. (D) The three stages are spread over the duration of the simulation with prescribed amplitudes (amp.). (E) Once equilibrated, we identify a specified number of chordal insertion sites using rejection sampling. (F) Next, we create “stiff” chordae in the end-systolic configuration and reverse leaflet deformations. (G) We then pressurize the valve with “stiff” chordae. (H) Next, we calibrate the end-diastolic length of each chord for an Ogden material law by identifying the ideal stretch value (*λ*^*∗*^) for the measured 1^*st*^ Piola-Kirchhoff stress (*P\**) in the corresponding “stiff” chord. (I) Finally, we pressurize the calibrated valve to obtain the final end-systolic configuration.

### Shape Field Refinement

Upon calibrating downward and lateral force values and pressurizing the valve leaflets against the rigid template, we obtain an initial result from our shape-matching framework. However, contact with the rigid template prevents the valve leaflets from equilibrating at our desired target shape. To refine shape-matching performance, we use a locally corrective pressure field (LCPF) in Stage II to penalize any deviations from the target shape, i.e., the end-systolic configuration, see **Fig. 3B**. To do so within our finite element simulations, we must define a tractable, analytical expression that reproduces the shortest vector distance from any point to our target surface. Here, we first determine the maximum and minimum dimensions of our target surface in the x, y, and z dimensions in MATLAB 2024b (MathWorks Inc., Natick, MA). We then extend this domain by 2 mm at each end. Next, we sample points at every 0.5 mm within this domain and determine the shortest vector to the target surface at each point using the point2trimesh package[28]. Thus, we generate a vector field for 74*×*66*×*50 points. We then fit a cosine series to our distance field with 732,600 and threshold the bottom 1% of frequencies to remove 92.21% of components. Unlike the other heart valves, the lack of anatomic symmetry in the tricuspid morphology leads to a reduction of components that only marginally improves computational tractability at this stage. To further compress the shape field around our target shape, we first apply a 3D Gaussian filter to the field to smooth out components. Further details on Gaussian smoothing are provided in the **Supplementary Materials**. We then proceed as before by fitting a cosine series to our smoothed field and thresholding the bottom 1% of frequencies to remove 99.77% of components. As a result, we decrease the number of active components by 33*×*; 57,072 for thresholding alone vs 1,695 for smoothing then thresholding. Through this process, we penalize any discrepancy in position between our finite element mesh and the target shape with a corrective pressure of 10 kPa per mm.

Finally, in Stage III we disable contact with the rigid template and allow the leaflets to equilibrate under the transvalvular pressure, CMF, and LCPF, see **Fig. 3C**. The (de)activation of each of the aforementioned forces and constraints is carefully controlled in Abaqus through user-specified linear and “smooth-step” amplitudes, see **Fig. 3D**.

#### Algorithm 1

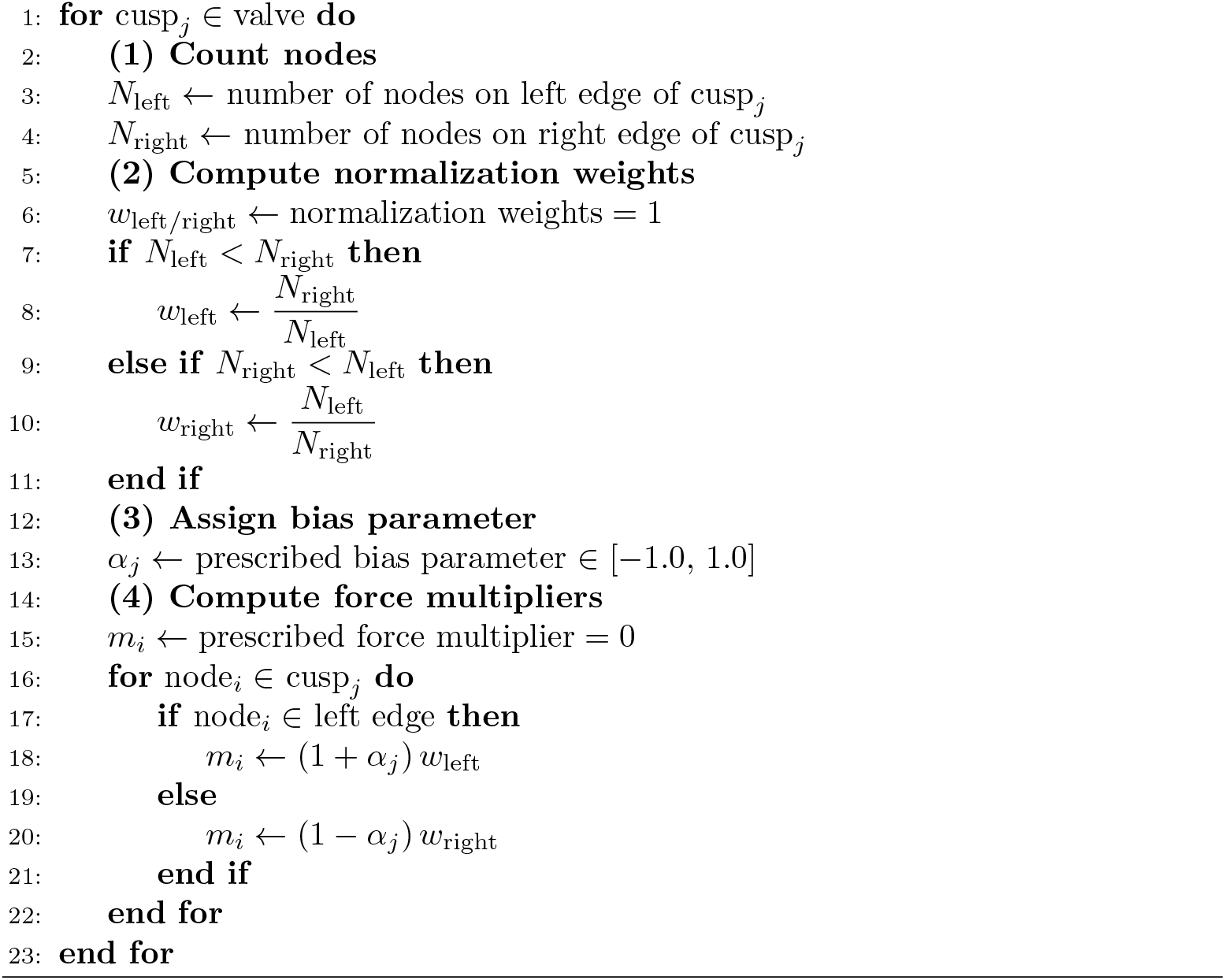

Computation of cusp bias and force multipliers

### Synthetic Chordal Insertions

#### Generating Insertion Locations

Once we establish a correspondence between the end-diastolic and end-systolic configurations of the valve, we create synthetic chordal insertions. Similar to Khalighi et al[23], we create individual, branchless chordae tendineae that produce an equivalent tethering effect to ground-truth insertions, or “functionally equivalent” chordae.

In clinical applications, the location of papillary muscle heads can be identified via non-invasive imaging and may be provided as inputs to our framework[29]. In this study, we first assign papillary muscle locations used in the original finite element model. Next, we randomly insert chordae into the ventricular surface of the valve in the deformed configuration. Specifically, we distribute 25% of chordal insertions in the buckling or fold zones of the leaflets and 25% of insertions along the free edge, see **Fig. 3E**. Furthermore, we remove insertions that are assigned to nodes along the valve annulus via rejection sampling. Next, chordal insertions are assigned to the nearest papillary muscle in the deformed configuration from a given insertion point. Finally, any chordae that will not buckle upon the reversal of deformation are removed.

#### Calibrating Chordal Lengths

We reverse deformations from the end-systolic to end-diastolic configuration with nearly-rigid or “stiff” chordae tendineae, see **Fig. 3F**. To that end, chordae tendineae are discretized using two T3D2 elements in Abaqus and we use a Neo-hookean material model with modulus 10 GPa. We then pressurize the valve with these “stiff” chordae to determine reaction forces at their base, see **Fig. 3G**. Next, we invert the stress-stretch relationship for chordae tendineae to determine the initial length of each chord, see **Fig. 3H**. Specifically, we identify a stretch of *λ*^*∗*^, which produces a 1^st^ Piola-Kirchhoff stress value of *P*^*∗*^ in a truss element using the Ogden model. Here, the Ogden material coefficients are obtained from prior in vitro uniaxial tensile testing of tricuspid chordae tendineae, as described above. Once calibrated, we pressurize the valve to 22.95 mmHg (as measured in the ex vivo beating heart[24]), to simulate leaflet coaptation with synthetic chordae, see **Fig. 3I**.

#### Sensitivity to Insertion Number

We also examine the sensitivity of valve coaptation to the number of synthetic chordal insertions. To do so, we identified the number of chordal insertions in eight explanted, structurally intact human tricuspid valves. Specifically, we marked insertion locations on images of the ventricular surface of each valve taken during planimetry using an in-house MATLAB script. This study was performed in a blinded manner, and its results are summarized in **Table 1**. Please note that we round down to the nearest natural number when assigning insertions based on count or (area) density in our simulations.

**Table 1.**
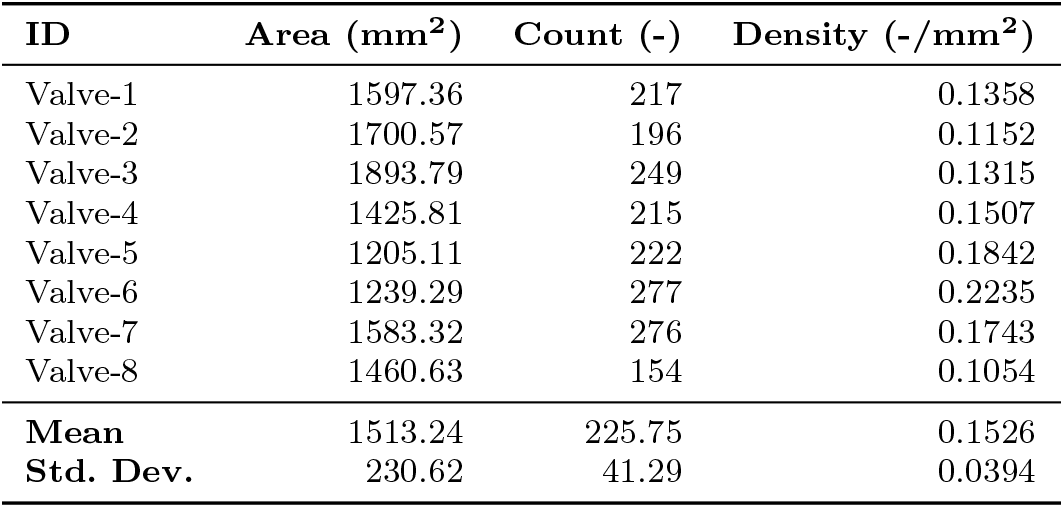
Chordal insertion density for explanted, structurally intact, human tricuspid valves (n=8) determined through a blinded, reader study.

#### Evaluation metrics

We evaluate our simulation results by computing functional and engineering metrics at end-systole. Functional metrics include the mean inter-surface distance, computed as described in **Equation 2**, and the leaflet coaptation area[30]. Engineering metrics include the maximum principal stretch (*λ*_1_), computed as *λ*_1_ = exp(*H*_1_), where *H*_1_ is the maximum principal Hencky strain output from Abaqus. We also compute the areal strain, which is defined as the relative change in element areas between the (end-diastolic) and deformed (end-systolic) configurations. All quantities are presented as mean *±* one standard deviation unless specified otherwise.

## 3 Results

### Model validation against beating heart echocardiography

To validate Texas TriValve 1.1, we imposed the end-systolic Neumann and Dirichlet boundary conditions and quasi-statically simulated full closure of the valve with the explicit finite element method. We then compared the 3D valve geometry to the 2D echocardiography data collected in the beating human heart. To this end, we first identified the echocardiographic imaging plane in our 3D model, see **Fig. 4A**. Next, we measured distances between key landmarks in both the echocardiographic images and in our 2D plane-cut of the 3D model. Specifically, we identified four landmarks: anterior leaflet hinge point (A), septal leaflet hinge point (S), point of coaptation between the two leaflets (C), and the coaptation point of the leaflet free edges (T), see **Fig. 4B**. Based on these measurements, we found that our model configuration closely matched our echocardiographic data: We found an S-A distance of 20.6 mm in both model and experiment, an S-C distance of 10.5 mm and 10.4 mm in model and experiment, respectively, an A-C distance of 12.6 mm and 11.3 mm again in model and experiment, respectively, and a C-T distance of 3.9 mm and 6.1 mm in the model and experiment, respectively. All discrepancies in measurements lie within the largest voxel size, 3 mm, of 3D echocardiographic probes[27].

**Fig 4.**
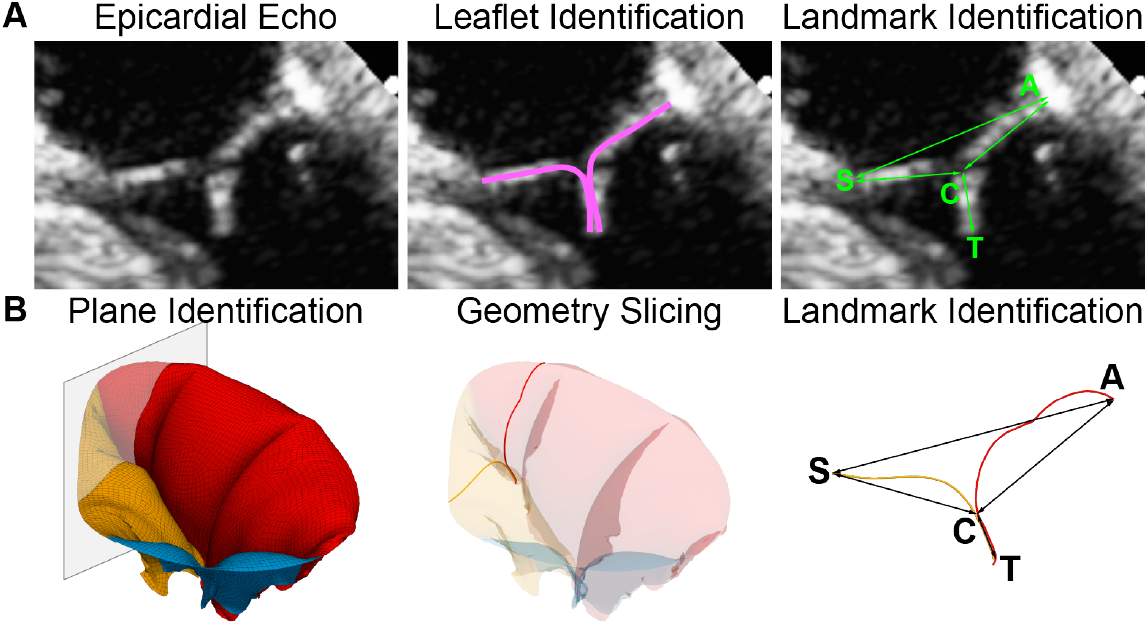
Texas TriValve 1.1 validation against beating heart echocardiography: (A) To validate our valve simulations we measured the distances between the anterior (A) and septal (S) leaflet hinge points as well the coaptation point (C) and coaptation length (T) in 2D echocardiographic images in the beating heart. (B) We then identified the corresponding imaging plane in our finite element model and repeated the same measurements as in the echocardiographic images for comparison.

### Shape matching recreates end-systolic deformations

Our tricuspid valve-specific, hyperelastic shape matching procedure produces an end-systolic configuration that closely matches the ground truth; see **Fig. 5A**. Upon examining cross-sectional slices between different cusp pairs in the valve, we can observe the progression of the shape matching process, **Fig. 5B&C**. In Stage I, pressurization against the rigid template and the application of downward and lateral forces ensure the correct combinations of cusps form contact pairs. Please note that the leaflets are still separated by the rigid template and are not in direct contact with one another. However, there are shape errors that appear in the leaflet centers or belly regions. These are largely eliminated with the addition of the pressure field in Stage II. Finally, removing the rigid template in Stage III allows the leaflets to establish contact with one another and energetically equilibrate. In certain cross-sectional planes, the predicted leaflet coaptation line moves above the target shape’s coaptation line in the equilibrated configuration, as seen in Slices #2, #4, and #5.

**Fig 5.**
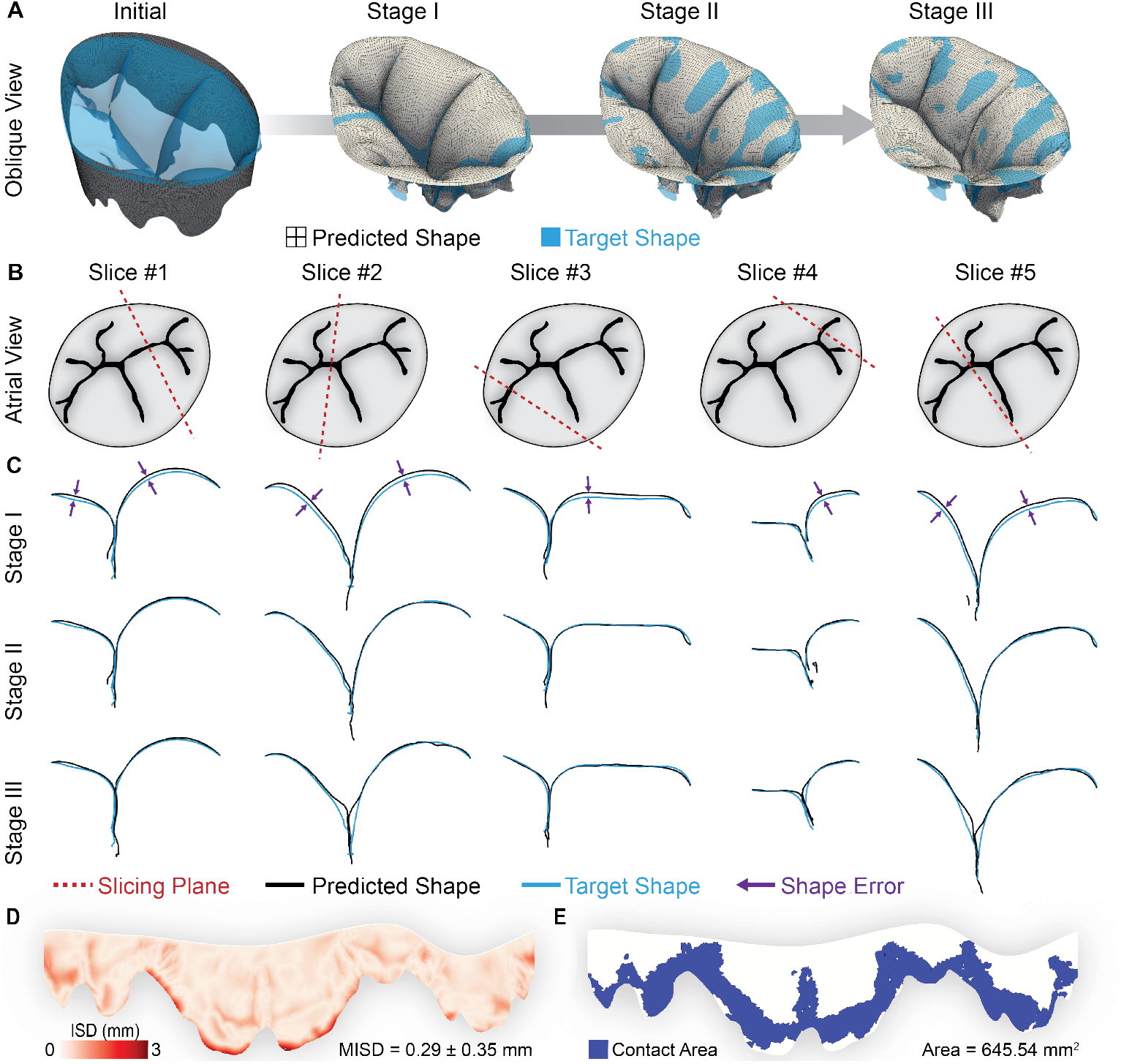
Shape matching recapitulates leaflet contact: (A) The multi-stage hyperelastic shape-matching process yields good agreement between the target and predicted shape. (B) We slice our simulation domain along five representative planes that feature different inter-leaflet contact pairs. (C) When examining individual slices, we see that pressurization against the rigid template (Stage I) provides coarse agreement in contact pairing between the predicted and target shapes. Next, errors between both shapes are minimized in the leaflet belly regions with the activation of the shape field (Stage II). Finally, removing the rigid template (Stage III) ensures that leaflets in the predicted shape coapt in their final state.

Upon quantitative comparison of leaflet deformations between our target and predicted shapes, we find a mean inter-surface distance (ISD) of 0.29 *±* 0.35 mm. ISD values cluster closer to 0 along the coaptation line and leaflet centers, with higher values observed at the free edge and a peak value of 3.03 mm, see **Fig. 5D**. Notably, the predicted shape overestimates leaflet contact area by 16.88% as compared to the target shape (645.54 mm^2^ vs 552.33 mm^2^), see **Fig. 5E**.

### Synthetic chordae recreate end-systolic deformations

We consider the equilibrated valve at the end of Stage III to be the end-systolic state and build synthetic chordae as described above. Furthermore, to determine the sensitivity of the predicted shape to the number of chordal insertions, we compare three distinct cases to the “Target” shape: (i) synthetic insertions based on measured area-density (n=202) or “Density”, (ii) synthetic insertions based on measured count (n=225) or “Count”, and (iii) synthetic insertions based on twice the measured count (n=450) or “2X Count”, see **Fig. 6A**.

**Fig 6.**
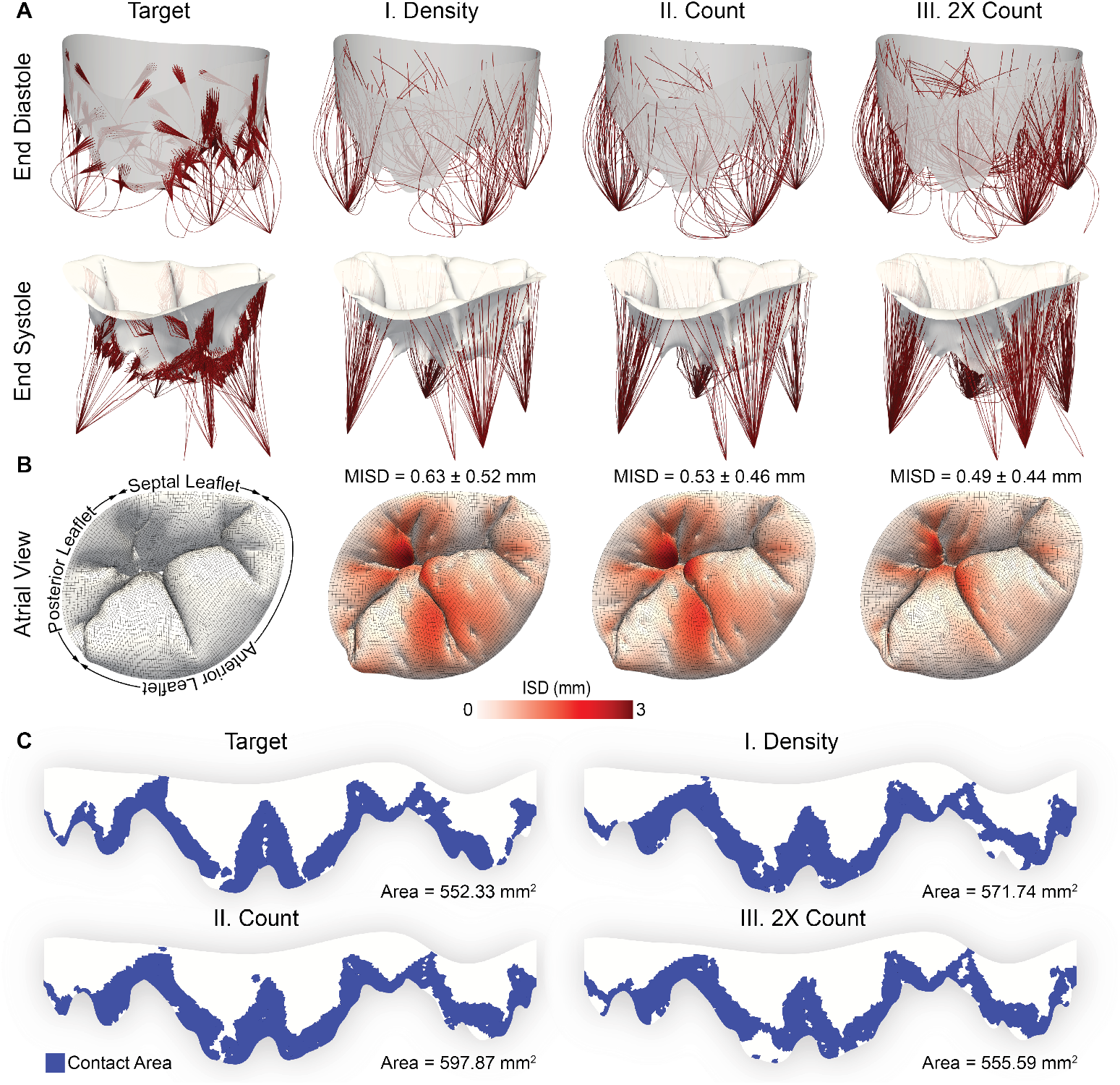
Tricuspid valve functional metrics are sensitive to the number of synthetic chordae: (A) Against the “Target”, we compare leaflet deformations at the end-systolic state for three distinct cases: Density, Count, and 2X Count. (B) Atrial view of different chordal configurations overlaid with contours of inter-surface distance (ISD) at end-systole. (C) Contact area projected onto a two-dimensional representation of the tricuspid valve leaflets at end-diastole. ISD presented as Mean *±* Std. Dev.

We see that insertions based on Density, Count, and 2X Count produce end-systolic configurations similar to the Target, see **Fig. 6B**. Notably, we observe that 2X Count almost exactly reproduces coaptation lines between the different tricuspid leaflets. Please note that the predicted shape presents a “pincushion” pattern due to localized leaflet stress. In the target shape, chordal insertions include a radial fan emerging from the main insertion that relieves any such stress localization. Quantitatively, we observe that increasing the number of chordae decreases the inter-surface distance (ISD) between the target and predicted shapes. Specifically, we note the mean ISD is 0.63 *±* 0.52 mm for Density, 0.54 *±* 0.46 mm for Count, and 0.49 *±* 0.44 mm for 2X Count. In comparison, increasing the number of chordae non-uniformly alters the contact area of each valve, see **Fig. 6C**. Here we see a 3.52% increase in area for Density (571.74 mm^2^), a larger 8.25% increase in area for Count (597.87 mm^2^), and a minimal increase of 0.59% in area for 2X Count (555.59 mm^2^).

### Synthetic chordae reproduce end-systolic leaflet stretch and strain

To determine the sensitivity of chordal insertion number to leaflet stretch and strain values, we average these quantities in the centers or “belly” regions of each leaflet where shape errors are minimal, see **Fig. 7A**.

**Fig 7.**
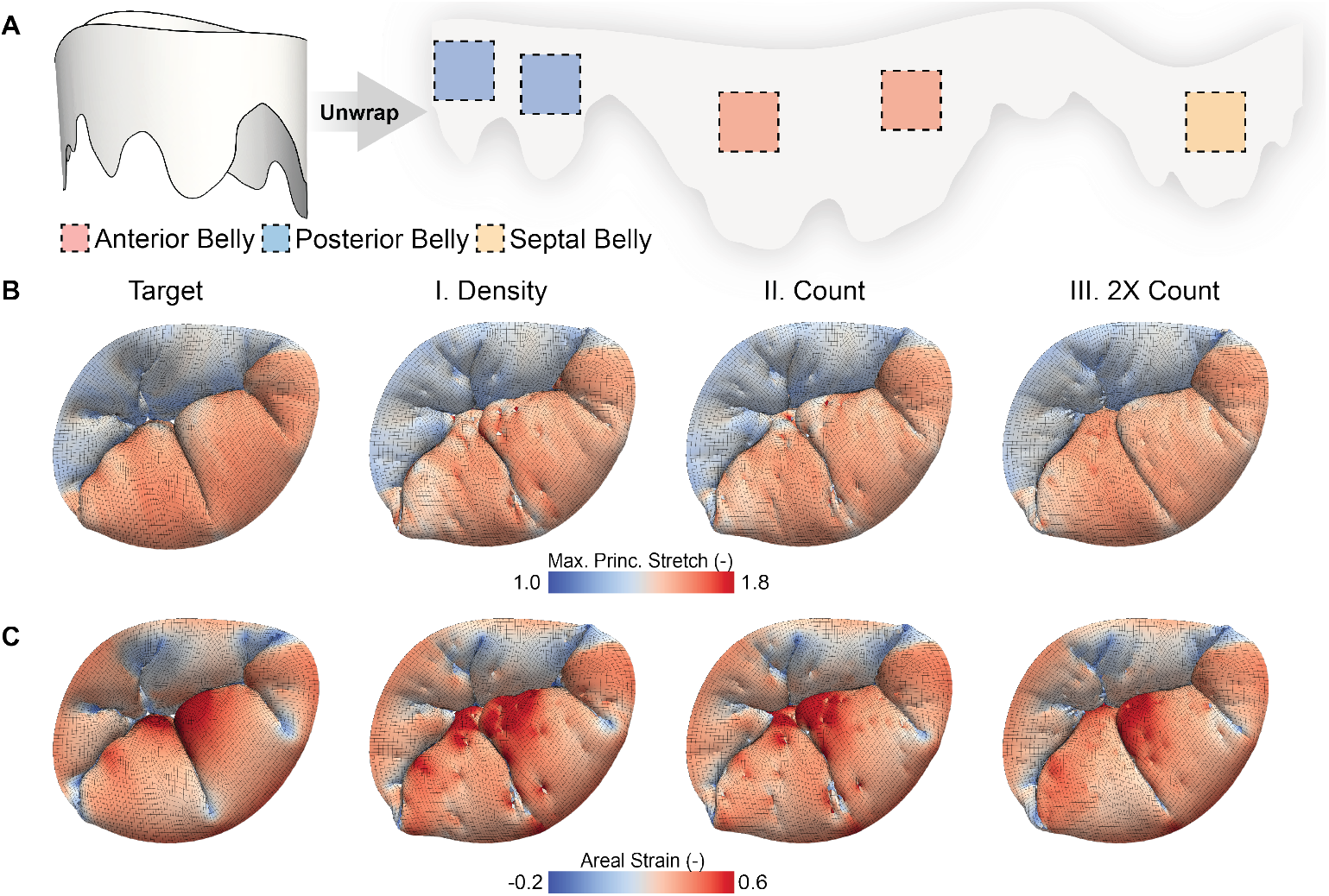
Tricuspid leaflet belly stretch and strain values are sensitive to the number of synthetic chordae: (A) We identify 7 mm × 7 mm regions in the center, or ‘belly’, of each leaflet on a 2D projection of the valve in the end-diastolic state. Contours of (B) maximum principal (max. princ.) stretch and (C) areal strain overlaid on an atrial view of the tricuspid valve at end-systole for each case.

Here, we found that all three configurations reliably recreate the spatial distribution of maximum principal stretch when compared to the target, see **Fig. 7B**. In detail, we observe that mean stretch values in the anterior leaflet belly are marginally underpredicted by approximately 2% in all cases. For the posterior leaflet belly, mean stretch values differ by less than 1% in all cases. Finally, mean stretch values differ by less than 2% in the septal leaflet belly. In summary, maximum principal stretch errors are limited for all configurations.

Similarly, we found that all three configurations broadly reproduce the spatial distribution of areal strain when compared to the target, see **Fig. 7C**. However, mean areal strains are underpredicted by 1.67-8.80% in the anterior leaflet belly, by up to 10.01% in the posterior leaflet belly, and by 3.38-4.07% in the septal leaflet belly. In summary, areal strain errors vary by leaflet across all three configurations. Stretch and strain values at the leaflet belly regions for the above cases are provided in **Tables 2** and **3**, respectively.

**Table 2.** Maximum principal stretch values in leaflet belly regions. All values are presented as mean *±* standard deviation.

| Chordal Set | Anterior Leaflet |  | Posterior Leaflet |  | Septal Leaflet |  |
| --- | --- | --- | --- | --- | --- | --- |
|  | Stretch(-) | Change(%) | Stretch(-) | Change(%) | Stretch(-) | Change(%) |
| Target | $1.60 \pm 0.03$ | – | $1.37 \pm 0.03$ | – | $1.38 \pm 0.04$ | – |
| Density | $1.56 \pm 0.04$ | –2.39 | $1.36 \pm 0.04$ | –0.97 | $1.36 \pm 0.04$ | –1.48 |
| Count | $1.56 \pm 0.04$ | –2.36 | $1.36 \pm 0.03$ | –0.87 | $1.36 \pm 0.03$ | –1.82 |
| 2X Count | $1.56 \pm 0.04$ | –2.00 | $1.37 \pm 0.04$ | –0.36 | $1.36 \pm 0.03$ | –1.54 |

**Table 3.**
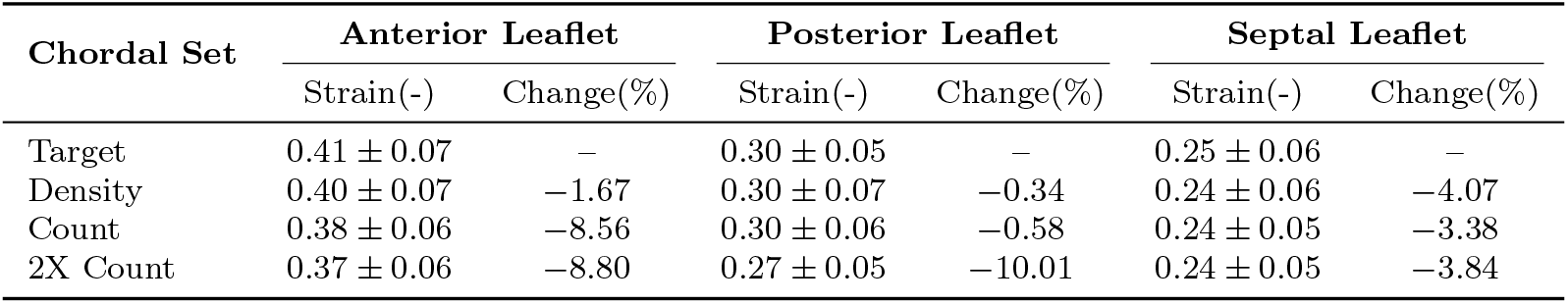
Areal strain values in leaflet belly regions. All values are presented as mean *±* standard deviation.

| Chordal Set | Anterior Leaflet |  | Posterior Leaflet |  | Septal Leaflet |  |
| --- | --- | --- | --- | --- | --- | --- |
|  | Strain(-) | Change(%) | Strain(-) | Change(%) | Strain(-) | Change(%) |
| Target | $0.41 \pm 0.07$ | — | $0.30 \pm 0.05$ | — | $0.25 \pm 0.06$ | — |
| Density | $0.40 \pm 0.07$ | -1.67 | $0.30 \pm 0.07$ | -0.34 | $0.24 \pm 0.06$ | -4.07 |
| Count | $0.38 \pm 0.06$ | -8.56 | $0.30 \pm 0.06$ | -0.58 | $0.24 \pm 0.05$ | -3.38 |
| 2X Count | $0.37 \pm 0.06$ | -8.80 | $0.27 \pm 0.05$ | -10.01 | $0.24 \pm 0.05$ | -3.84 |

## 4 Discussion

A critical challenge in creating predictive models of the tricuspid valve is our inability to characterize chordal attachments through non-invasive imaging. In this study, we propose novel methods to overcome this by creating synthetic chordal attachments for tricuspid valve models. To do so, we first modify an established shape matching technique with anatomic insights from explanted tricuspid valves and improve its computational tractability through Gaussian smoothing. We then use zone-based rejection sampling to seed chordal insertions and a stress-based calibration process to create synthetic chordae. Finally, we examine the sensitivity of functional and engineering metrics to the number of chordal insertions. Together, these activities allow us to achieve our stated objective of predicting tricuspid valve end-systolic deformations *without* access to, or knowledge of, in vivo subject-specific chordal geometry.

The synthetic chordal insertions tether the valve leaflets and produce deformations that closely agree with our target shape: the finite element model of a reverse-engineered human tricuspid valve in the end-systolic configuration. To that end, they yield mean inter-surface distance errors well below 1 mm and errors in contact area between 0.59%–8.25%. They also yield errors in maximum principal stretch between 0.36%–2.39% and errors in areal strain between 0.34%–10.01% in leaflet belly regions across all cases. Based on these figures, for our exemplar valve we recommend n=450 chordal insertions to minimize errors in distance, contact area, and maximum principal stretch. Furthermore, our results are comparable to, and in some cases outperform, existing shape matching or chordae generation methods in the literature. For example, Wu and colleagues use an LDDM-like method to predict atrioventricular valve strains[31]. They obtain mean absolute errors of 0.138 and 0.055 in areal and maximum principal strains, respectively, across mitral leaflets in a benchmark problem. In comparison, our synthetic insertions produce mean absolute errors between 0.076–0.085 and 0.245–0.287 in areal and maximum principal strains across the entire tricuspid valve, respectively. Please note that we convert our model’s maximum principal stretch values to their corresponding Green-Lagrange strain values here. Similarly, Rego et al. obtained stretch residuals of 0.006 *±* 0.115 in the circumferential direction and *−*0.009 *±* 0.131 in the radial direction using hyperelastic shape matching[20]. Notably, we observed maximum principal stretch residuals between 0.000 ± 0.158 and 0.006 *±* 0.181 in our simulations. Finally, we compare our results with two prior studies that create synthetic chordae and report inter-surface distances or *l*^*2*^ errors. First, Kong et al. create synthetic chordae for imaging-based tricuspid valve models and iteratively adjust chordal lengths to obtain mean *l*^*2*^ errors between 0.55-0.60 mm, with a stated objective of ensuring leaflet belly errors below 2 mm. Second, the work of Liu et al. is closest to our current work as they use both hyperelastic shape matching and stress-based calibrations, alongside the addition of leaflet and chordal pre-stretch, to create synthetic chordal insertions. Specifically, they report mean *l*^*2*^ errors between 0.38-0.58 mm, with a stated objective of ensuring mean errors below 1 mm. Both results are comparable to our own mean *l*^*2*^ errors that lie between 0.49– 0.63 mm, and are well under the resolution of 3D echocardiographic probes[27]. The above quantitative comparisons are summarized in **Table A1**.

The potential impact of our work is twofold. Firstly, we develop and open-source a novel shape matching algorithm for human tricuspid valves. In particular, we incorporate anatomic insights from explanted tricuspid valves to accommodate the lateral tension exerted by tricuspid chordae tendineae on valve leaflets[26]. By using a Gaussian filter, we also reduce the computational cost of DCT-based hyperelastic shape matching for the irregular tricuspid valve morphology. Unlike the consistent bi-leaflet morphology of the mitral valve[32], Hahn and colleagues describe six distinct variations in tricuspid leaflet morphology in an echocardiographic study[33]. Furthermore, we posit that our framework is robust to such morphological variations, as demonstrated by its application to a rare Hahn-Type IV valve (incidence of 2%) with five cusps. Nevertheless, further testing on other morphologically diverse subjects is warranted. Secondly, we perform *the first* sensitivity study of predicted valvular function to chordal insertion number for a human tricuspid valve model. Prior studies have used an insertion density of 0.15–0.18 insertions/mm^2^ to model the mitral valve [23, 34, 35]. When applied to our exemplar valve, this would produce between 198–238 chordae, and is similar to our density- (n=202) and count-based (n=225) insertions. Our results indicate that more than twice this number of chordal insertions (n=450) are needed to accurately reproduce deformations in our exemplar tricuspid valve. Interestingly, a prior study has shown that the tricuspid valve is tethered by up to twice as many chordae as the mitral valve across three distinct large-animal models[36]. Thus, there may be an anatomic foundation to our results that must be investigated more deeply.

Naturally, our study is subject to several limitations. Firstly, we use leaflet- and chordae-specific material properties determined from in vitro mechanical characterization. In clinical use, such information is unavailable, and inverse methods may be needed to identify patient-specific material properties[37]. Secondly, we currently create branchless, synthetic chordae that are functionally equivalent to their patient-specific counterparts. However, recent studies indicate that branching parameters may affect leaflet coaptation areas and must be considered in future studies[38]. Thirdly, we demonstrate our pipeline using a tricuspid valve geometry reconstructed from annular measurements in an ex vivo beating heart and through in vitro morphological characterization. Future studies will use imaging-derived leaflet geometries, and we anticipate no challenges in applying this framework there. Finally, we manually assign bias parameters to the lateral chordal mimicking forces due to the significant computational cost of simultaneously optimizing six variables[39]. Moving forward, we will overcome this challenge by optimizing bias parameters independently for each leaflet in a sequential manner.

## 5 Conclusion

In conclusion, we developed a novel framework to predict tricuspid valve coaptation without explicit in vivo chordal geometry. To do so, we combined an updated hyperelastic shape matching framework with a stress-based calibration of functionally equivalent chordae tendineae. We demonstrate the utility of this approach by predicting end-systolic deformations in a morphologically rare, reverse-engineered model of the human tricuspid valve, Texas TriValve 1.1. In addition, we examine the sensitivity of valvular function to the number of chordal insertions and provide preliminary recommendations for insertion count. Notably, we open-source our models and subroutines so that others may implement similar features within their own modeling frameworks. We envision this work as taking the first step towards establishing morphologically-varied guidelines for synthetic chordal generation. Such guidelines may enable the large-scale use of predictive tricuspid valve models in clinical applications which, in turn, may improve patient outcomes.

## Acknowledgements

This study was supported, in part, by awards from the National Heart, Lung, and Blood Institute to MKR (R01HL165251, R21HL161832, R21HL170177) and MSS (R01HL184128), and through a predoctoral fellowship to CEH (F31HL178280) and NTS (F31HL170754). This work was also supported, in part, by the American Heart Association through an award to WH (25IPA1454136) and a predoctoral fellowship to MM (902502).

## Data Availability

Texas TriValve 1.1 is also openly available on GitHub: https://github.com/SoftTissueBiomechanicsLab/Texas_TriValve_1.1. Moreover, Abaqus input files and user subroutines for hyperelastic shape matching and synthetic chordal insertion are also available on GitHub: https://github.com/SoftTissueBiomechanicsLab/No_Strings_Attached.

## Conflicts of Interest

MKR has a speaking agreement with Edwards Lifesciences and LumenexBio.

**Table A1.** Quantitative comparison to selected prior studies with atrioventricular shape matching or synthetic chordal insertions. Summary statistics are presented as mean *±* 1 standard deviation, while ranges are presented as [Lower Bound, Upper Bound] where applicable. Circ.: Circumferential; Rad.: Radial; MAE: Mean Absolute Error; *: Maximum (Max.) Principal (Princ.) Stretch Values.

| Prior Study | Valve | Metric | Reported Value | Our Study |
| --- | --- | --- | --- | --- |
| *Shape Matching* |  |  |  |  |
| Wu et al.[31] | Mitral | Areal Strain MAE (-) | 0.138 | [0.076,0.085] |
|  |  | Max. Princ. Strain MAE (-) | 0.055 | [0.245, 0.287] |
| Rego et al[20] | Mitral | Circ. Stretch Residual (-) | $0.006 \pm 0.115$ | $[0.000 \pm 0.158,$ |
| | | Rad. Stretch Residual (-) | $-0.009 \pm 0.131$ | $0.006 \pm 0.181]^*$ |
| *Synthetic Chordae Creation* |  |  |  |  |
| Kong et al.[40] | Tricuspid | Mean $l^2$ Error (mm) | [0.55,0.60] | [0.49,0.63] |
| Liu et al.[35] | Mitral | Mean $l^2$ Error (mm) | [0.38,0.58] | [0.49,0.63] |

