## Supplemental_Materials for "No Strings Attached: Predicting Tricuspid Valve Deformation Without In Vivo Chordal Geometry"

### S1 Effect of Lateral Chordal Mimicking Force

In this study, we use lateral chordal mimicking forces to ensure that there is adequate contact between cusps of the same tricuspid valve leaflet. In their absence, the intra-leaflet buckling zones are insufficiently resolved, see **Fig. S1**. While the lack of lateral CMFs doesn't seem to affect the atrial surface of the predicted shape, we observe large discrepancies in intra-leaflet contact for the anterior tricuspid leaflet.

### S2 Chordal Mimicking Force Calibration

As described in in the main text, we created both downward and lateral forces on the free edge of the leaflets to capture their tethering behavior in our hyperelastic shape matching procedure. Given that the total magnitude of these forces is unknown from 3D transesophageal echocardiography measurements, we first aimed to understand the amount of force that tethered the leaflets without hindering closure. We used Gaussian-process (GP) based Bayesian optimization to search for the downward and lateral chordal mimicking force (CMF) magnitudes that minimized the mean inter-surface distance between the simulation and the reference surface. To this end, we used MATLAB's `bayesopt` function to iteratively select combinations of downward and lateral force magnitudes. The scalar objective was defined as the mean inter-surface distance (ISD) between the respective CMF simulation and the reference surface, as given in **Equation (1)** of the main text. Simulations that failed to converge or otherwise did not return a finite error returned `NaN` and were treated as failed objective evaluations. Because identical CMF magnitudes produced identical Abaqus simulations, the objective function was specified as deterministic (`IsObjectiveDeterministic=true`). MATLAB's default expected-improvement-per-second-plus acquisition function and exploration ratio of 0.5 were retained, and simulations were evaluated using four parallel workers. The CMF magnitudes were defined as continuous log-transformed variables with bounds of [0.01, 2.0] N. We terminated the optimization after 108 objective evaluations, after which we selected the force combination corresponding to the minimum estimated mean ISD as the calibrated CMF magnitudes. The estimated objective surface of the GP is shown in **Fig. S2**. We see a relatively broad region of similarly low error for different combinations of CMF force magnitudes. The optimum downward and lateral CMF magnitudes were found to be 1.04 N and 0.44 N, respectively, which resulted in a mean ISD of 0.2948 mm. We rounded these values to 1.0 N and 0.4 N to avoid implying unwarranted precision and to make the results more general, which produced a small increase in the mean ISD to 0.2976 mm. These results demonstrate that the CMF plays an important role in stabilizing the simulation during our shape matching procedure. This also suggests that the shape matching procedure is relatively insensitive to the precise CMF magnitude, and that our procedure can work well even if a magnitude within the low-error region is used. We highlight this advantage as future studies can leverage these results to apply similar CMF magnitudes without having to re-optimize for a valve-specific magnitude. However, we note that these results were derived from our one valve; the suitability of these magnitudes for other tricuspid valve geometries remains to be established.

### S3 Pressure Field Sensitivity to Gaussian Smoothing

To decrease the computational cost, we smooth components of the hyperelastic shape field using a three-dimensional Gaussian filter, see **Fig. S3**. We also ran a sensitivity study on the Gaussian smoothing parameters used in the definition of the locally corrective pressure. These included the

Gaussian window size and the Gaussian window standard deviation. After calibrating the chordal mimicking forces, we parametrically varied these by first holding the window standard deviation at 2.25 mm and varying the window size from 3 to 19 mm as shown in **Fig. S4A**. Then, we held the window size at 11 mm and varied the window standard deviation from 0.75 to 3.25 mm as shown in **Fig. S4B**. We selected 5 mm for the window size and 1.25 mm for the window standard deviation as a balance between accuracy in the mean inter-surface distance and total run time of the simulation.

### Supplementary Figures

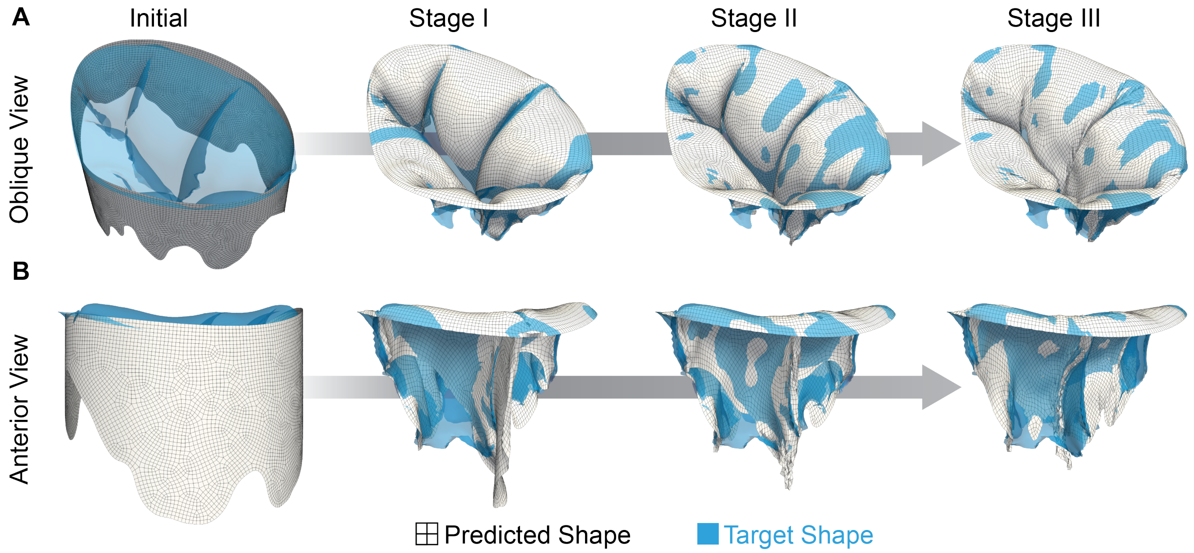


**Fig. S1: Coaptation Collapse Without Lateral CMF:** Shape matching procedure without application of lateral Chordal Mimicking Force (CMF) as observed from an (A) oblique view and (B) anterior leaflet-focused view. Here we see that the anterior leaflet folds up without lateral forces, as opposed to the target shape where the leaflets inflate and coapt.

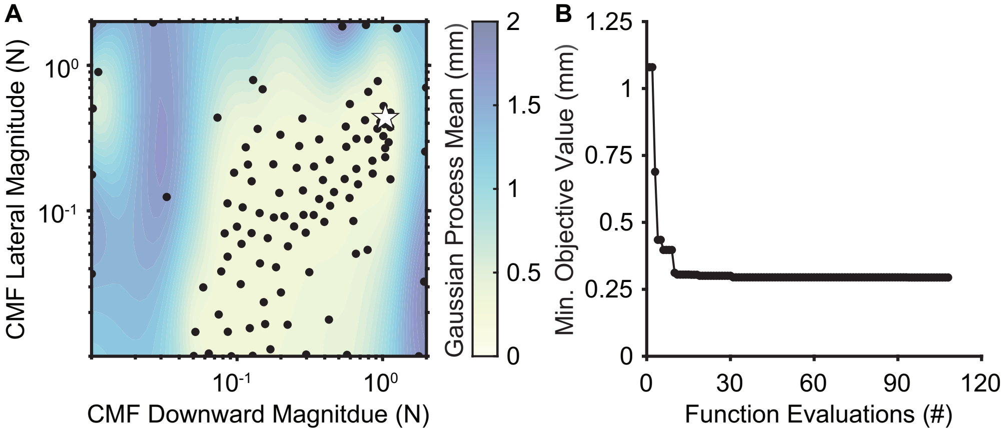


**Fig. S2: Sensitivity to Chordal Mimicking Force (CMF) Parameters:** (A) Individual function evaluations (black circles) are overlaid on contours of the mean response surface from the Gaussian process regression. The minimum observed value (white star) lies in a relatively flat region of the response surface. (B) Minimum (Min.) observed value of the objective function is obtained within 31 iterations and does not vary with additional evaluations. The optimal parameter value was found to be 1.04 N for the downward and 0.44 N for the lateral directions, with an error of 0.2948 mm.

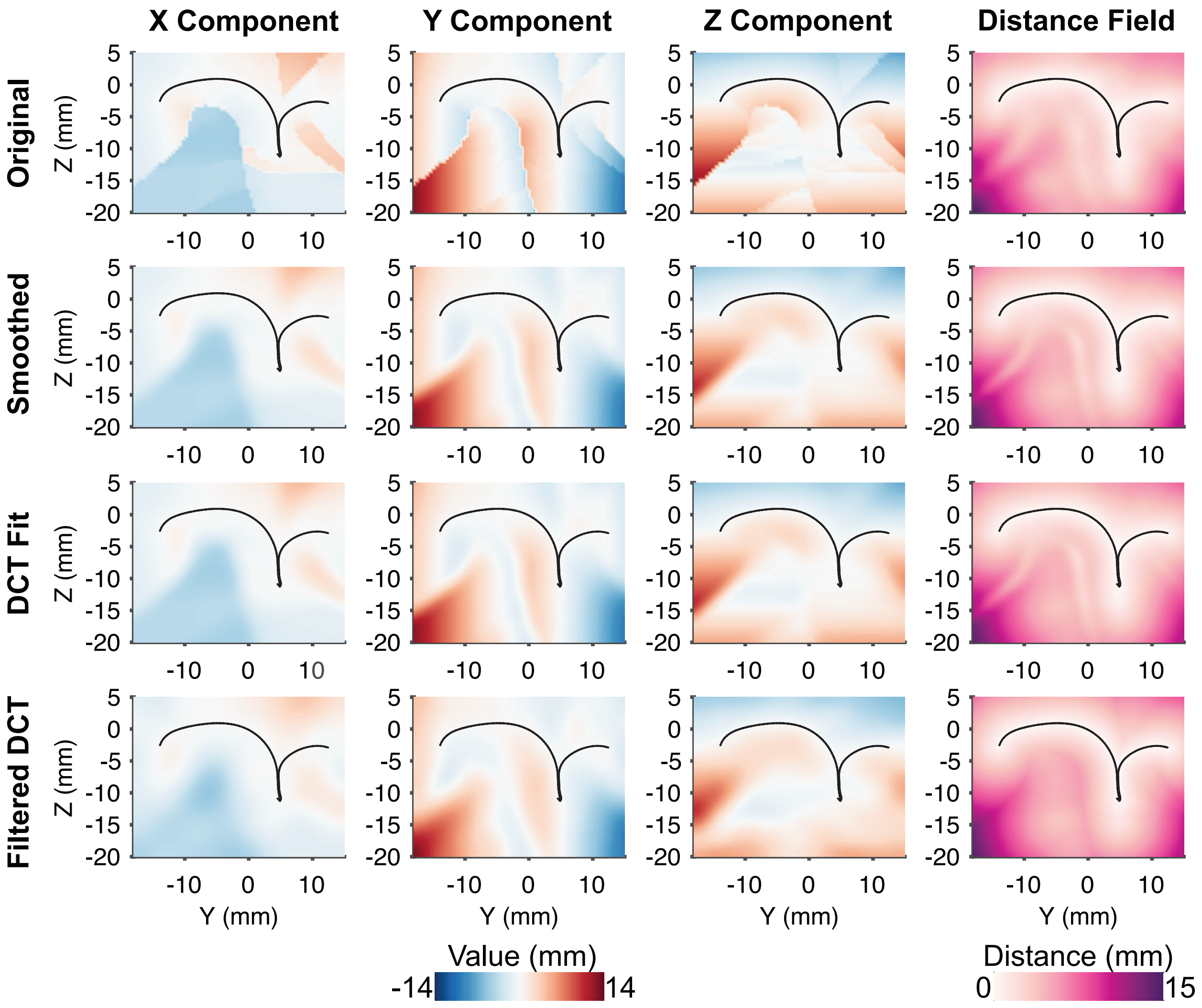


**Fig. S3: Compressing DCT Field:** We first smooth components of the nearest-point vector in the X, Y, and Z directions using a three-dimensional Gaussian filter. We then use a discrete cosine transform (DCT) to obtain an analytical representation of each smoothed vector field. Finally, we threshold 1% of the signal from each DCT fit to obtain a computationally tractable distance field for shape matching. The smoothing and filtering operations retain major patterns of the distance field, as seen in the cross-sectional views.

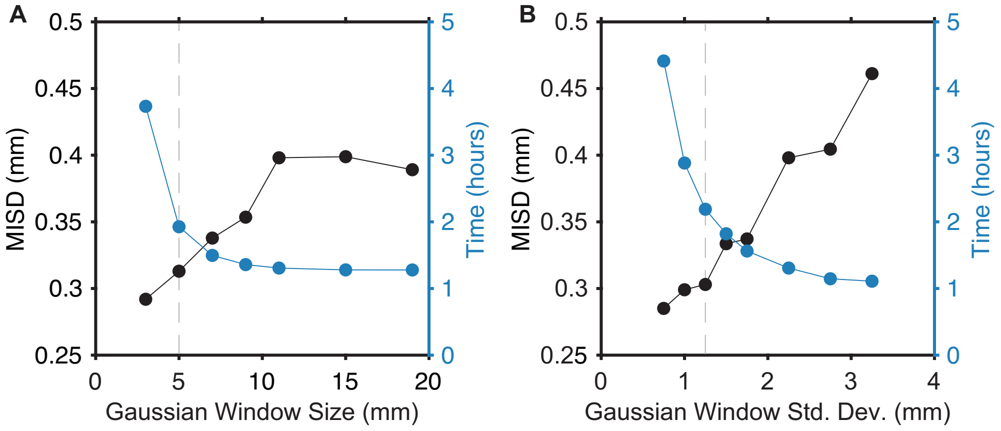


**Fig. S4: Sensitivity to Gaussian Smoothing Parameters:** We identify suitable parameter values for the Gaussian smoothing filter by balancing the Mean Inter-surface Distance (MISD) and Run Time for each parameter value. (A) For Gaussian window size, we vary parameter values between 3 mm and 19 mm. Similarly, (B) we identify a suitable Gaussian standard deviation (std. dev.) by varying values between 0.75 and 3.25. Vertical dashed lines indicate the selected parameter values, which are a window size of 5 mm and a standard deviation of 1.25 mm.
